# The cohesin modifier ESCO2 is stable during DNA replication

**DOI:** 10.1101/2022.06.21.497067

**Authors:** Allison M. Jevitt, Brooke D. Rankin, Jingrong Chen, Susannah Rankin

## Abstract

Cohesion between sister chromatids by the cohesin protein complex ensures accurate chromosome segregation and enables recombinational DNA repair. Sister chromatid cohesion is promoted by acetylation of the SMC3 subunit of cohesin by the ESCO2 acetyltransferase, inhibiting cohesin release from chromatin. The interaction of ESCO2 with the DNA replication machinery, in part through PCNA-interacting protein (PIP) motifs in ESCO2, is required for full cohesion establishment. Recent reports have suggested that Cul4-dependent degradation regulates the level of ESCO2 protein during late S/G2 phase of the cell cycle. To follow up on these observations, we have characterized ESCO2 stability in *Xenopus* egg extracts, a cell-free system that recapitulates cohesion establishment *in vitro*. We found that ESCO2 was stable during DNA replication in this system. Indeed, further challenging the system by inducing DNA damage signaling or increasing the number of nuclei undergoing DNA replication had no significant impact on the stability of ESCO2. In transgenic somatic cell lines, we did not see evidence of GFP-ESCO2 degradation during S phase of the cell cycle using either flow cytometry or live-cell imaging techniques. We conclude that ESCO2 is stable during DNA replication in both embryonic and somatic cells.

## Introduction

The tethering together of sister chromatids during DNA replication depends in part on acetylation of the SMC3 subunit of cohesin, which renders the complex resistant to removal from chromatin by the WAPL protein (1–3). In vertebrates, SMC3 acetylation is achieved by one of two related acetyltransferase enzymes, ESCO1 and ESCO2 (4). Using the *Xenopus* egg extract system, we previously showed that ESCO1 is developmentally regulated and not present at functional levels until after zygotic transcription begins (5). In egg extracts, therefore, ESCO2 is the sole cohesin acetyltransferase required for cohesion between sister chromatids, and depletion of ESCO2 from egg extract results in significant loss of cohesion (5, 6).

Multiple reports suggest cell cycle-dependent fluctuations in ESCO2 protein levels, although there are conflicting reports about the precise timing. Some reports indicate that ESCO2 levels peak during S phase (7)and are thus low prior to mitotic entry, while others have suggested that ESCO2 is degraded during M phase (4, 8). ESCO2 has also been reported to be stabilized by interaction with the MCM helicase during replication licensing, suggesting a third, perhaps indirect, level of stability control (9, 10).

ESCO2 protein levels are controlled at least in part by ubiquitin-dependent proteolysis (5, 8). The anaphase-promoting complex (APC) is an E3 ubiquitin ligase that has a number of substrates, including some that are degraded at mitotic exit, and others that continue to be recognized through G1 (11). As in other APC targets, a degron sequence in ESCO2 mediates recognition and modification by the APC when it is bound to the G1 specificity factor called Cdh1 (5, 11, 12). Mutation of this sequence stabilizes ESCO2, preventing its degradation in the presence of Cdh1 (5).

It has been suggested that degradation of ESCO2 is also controlled by a second E3 ubiquitin ligase, the Cul4-DDB1 complex via the specificity factor DCAF1 (Ddb1 and Cul4 associated factor 1, also called VprBP), resulting in degradation during late S and G2 (7). Together, these reports suggest an interesting dual regulation of ESCO2 by proteolysis: in G1 by the APC, and during S phase by the Cul4-DDB1-DCAF1^VprBP^ complex.

To better understand the regulation of ESCO2 protein turnover, we set out to identify the degron that might mediate recognition of ESCO2 by CUL4-DCAF1^VprBP^. To this end we analyzed ESCO2 stability, utilizing the *Xenopus* egg extract system, which is a powerful tool to investigate CUL4-dependent mechanisms (13–17). Our results indicate that ESCO2 is stable during DNA replication in the egg extract system. We also tested ESCO2 stability in cultured somatic cells, where we saw no evidence of degradation after G1 phase of the cell cycle. Our data suggest that accumulation of ESCO2 in the absence of CUL4-DCAF1^VprBP^ seen previously (7) may be developmentally regulated, a cell-type-specific response, or occur through indirect mechanisms.

## Results

Extracts prepared from the eggs of the frog *Xenopus laevis* are stockpiled with sufficient proteins for the replication of thousands of nuclei per microliter, making this system ideal for the study of DNA replication-dependent events *in vitro* (18, 19). Demembranated sperm heads added to the extract are assembled into nuclei through the recruitment of membrane vesicles from the extract and the import of nuclear and chromatin proteins. Regulated DNA replication and replication-dependent events such as cohesion establishment and Cul4-dependent degradation all occur in these *in vitro* assembled nuclei (6, 14, 15, 17).

To characterize the impact of DNA replication on ESCO2 stability we analyzed the level of ESCO2 protein in egg extract over time in the presence or absence of nuclei. Extracts were induced to enter interphase, then supplemented with nuclei (2300/μl), and the level of endogenous ESCO2 was assessed over time by western blot (Fig. 1A). In this experiment, ESCO2 levels were stable over the course of the experiment, and not impacted by the presence of nuclei in the extract (Fig. 1B). In contrast, the replication licensing factor Cdt1, a previously-characterized Cul4 substrate, showed clear nuclei-dependent degradation in the same extract. In control samples without nuclei Cdt1 levels decreased only slightly over the two-hour experiment, as previously reported, while in the presence of nuclei, Cdt1 was largely depleted by 60 minutes (14). Although there is virtually no transcriptional activity in egg extract (20), to rule out the possibility that the new translation of ESCO2 from maternal mRNA might mask our ability to detect protein loss, we added cycloheximide to the extract to block translation, and found that ESCO2 was unaffected (Fig. 2B). We conclude from this experiment that ESCO2 is stable during DNA replication in egg extract, though the Cul4-dependent degradation machinery is active.

**Figure 1.**
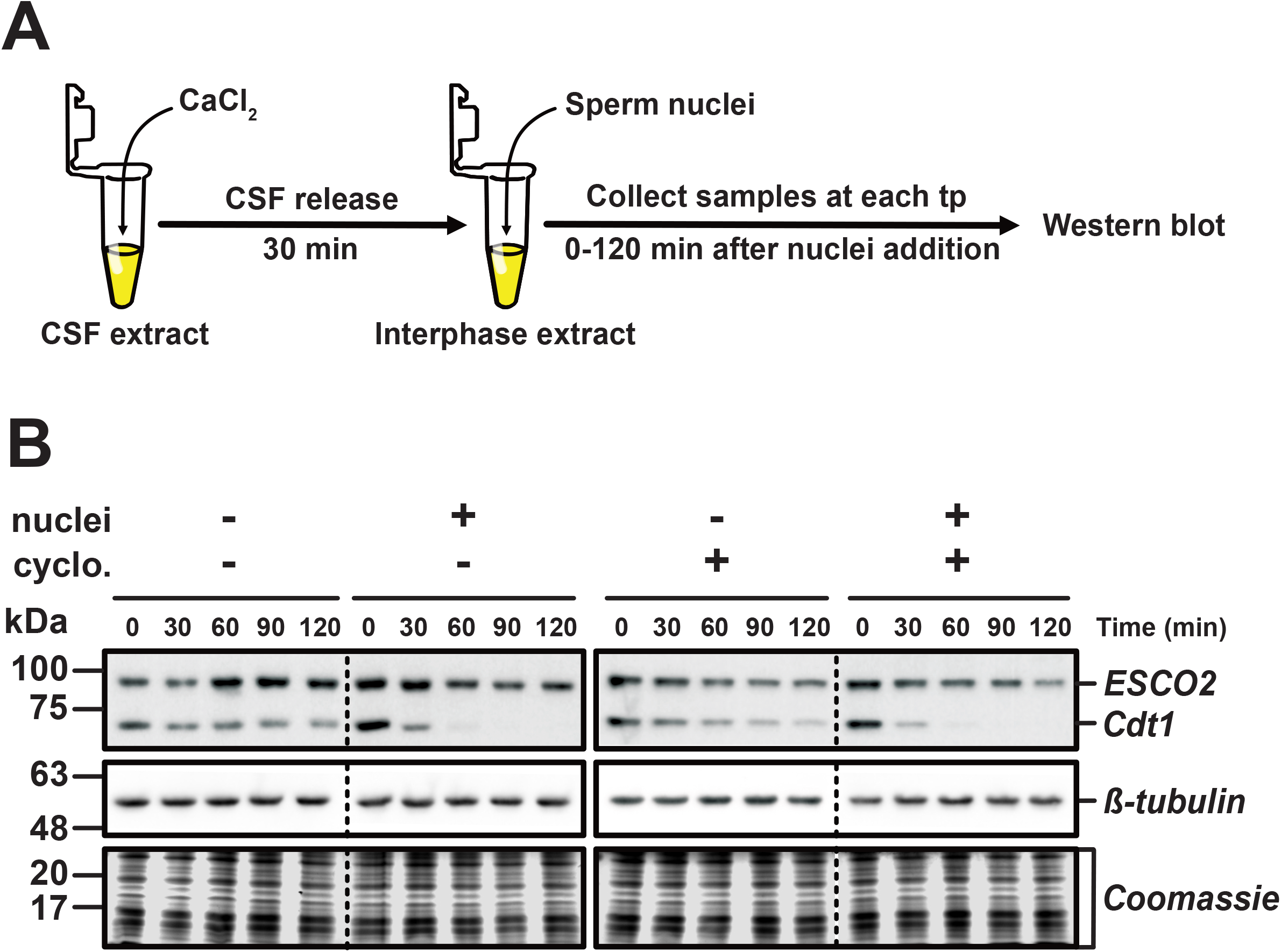
ESCO2 protein level remains constant during DNA replication. **A. Approach.** CSF-arrested extract was induced to enter interphase by the addition of calcium (CaCl_2_). Thirty minutes later, sperm nuclei (2300/μl, final) were added and aliquots were collected at the indicated times. **B. Immunoblot analysis.** Reaction samples were analyzed by immunoblot for the indicated proteins. **Cyclo:** Cyclohexamide was added where indicated to prevent protein translation. Solid outlines denote membrane fragments that were processed separately. Dotted lines denote where gel images were cropped. ESCO2 and Cdt1 were analyzed on the same membrane fragment. *β*-tubulin served as a loading control. The foot of the gel was collected and stained with Coomassie as an additional loading control. One representative experiment of three biological replicates is shown.

**Figure 2.**
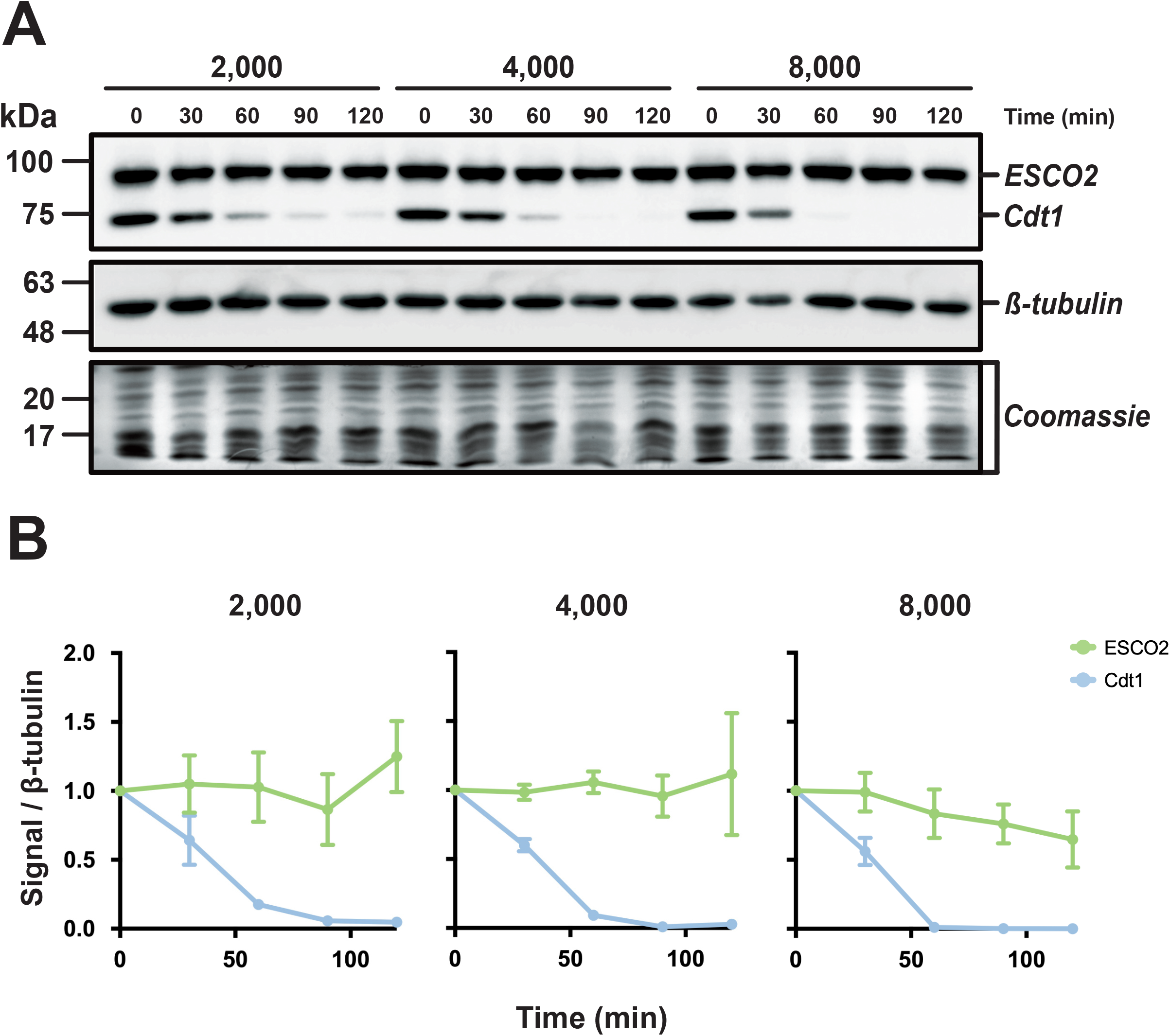
ESCO2 stability is unaffected by changes in the nucleus:cytoplasm ratio. **A. Immunoblot analysis.** Interphase extract was supplemented with the indicated concentrations of sperm nuclei (2000, 4000, or 8000 sperm nuclei/μl). Samples were collected at the indicated time points after the addition of sperm nuclei and were processed for immunoblot as in Figure 1B. Solid outlines denote membrane fragments that were processed separately. **B.** Quantification of results. The representative experiment shown in A, was repeated three times and the results were plotted as a fraction of the remaining signal for the indicated proteins normalized to the *β*-tubulin signal for each sample. Error bars = SD.

Although we saw no clear decrease in ESCO2 level during DNA replication in the egg extracts, we wondered whether ESCO2 stability might still be sensitive to the density of DNA replication, or the number of nuclei in the extract, perhaps being more efficiently degraded in the presence of increased nuclear density. To test this, we performed a titration experiment in which extract was supplemented with increasing concentrations of nuclei and analyzed the level of ESCO2 protein over time (Fig. 2). We found that increasing the nuclei concentration had no significant impact on ESCO2 protein levels. In fact, increasing the concentration of nuclei well above the highest nucleus:cytoplasm (N/C) ratio found in embryos following induction of zygotic transcription, estimated to be ~4000 nuclei/μl (20), had no impact on ESCO2 stability. Even at 8000 nuclei/μl ESCO2 levels were stable over the course of the experiment. In contrast, the degradation of Cdt1 was easily detectable at all nuclear densities, and slightly enhanced by the presence of additional nuclei in the extract (Fig. 2B). In the presence of 2000 nuclei/μl degradation was essentially complete by 90 minutes, while in the presence of 8000 nuclei/μl Cdt1 was undetectable at 60 minutes. We conclude from this experiment that ESCO2 degradation cannot be stimulated by increasing the N/C ratio, although Cdt1 degradation was enhanced under these conditions.

In budding yeast, the stability of the ESCO2 ortholog Eco1p is modulated by DNA damage signaling (21). To determine whether vertebrate ESCO2 stability might be regulated in response to DNA damage signaling, we tested two conditions known to activate the DNA damage response in egg extracts (Fig. 3A). First, we added the DNA polymerase inhibitor aphidicolin to the reaction. This drug causes the uncoupling of polymerase □ from the replicative helicase, resulting in the formation of single-stranded DNA, and a resultant DNA damage response (22–24). We also generated a DNA damage response by adding UV-irradiated sperm to the reaction (25). In both conditions, DNA damage signaling was verified by monitoring the phosphorylation of Chk1 checkpoint kinase (26) (Fig. 3A). Under these conditions, ESCO2 levels remained stable, while Cdt1 degradation seen previously was largely unaffected by whether the added sperm nuclei had been subjected to UV damage. Cdt1 degradation was partially inhibited by the presence of aphidicolin, consistent with the requirement for ongoing DNA replication for its degradation as shown previously (17). Consistent with this observation, inhibition of DNA replication by the addition of the Cdk inhibitor p27 also slowed Cdt1 degradation (Fig. 3A, right). We conclude from this experiment that ESCO2 stability is largely unaffected by DNA damage signaling, whether or not DNA replication is active, consistent with the notion that ESCO2 and Cdt1 stability are controlled by different mechanisms.

**Figure 3.**
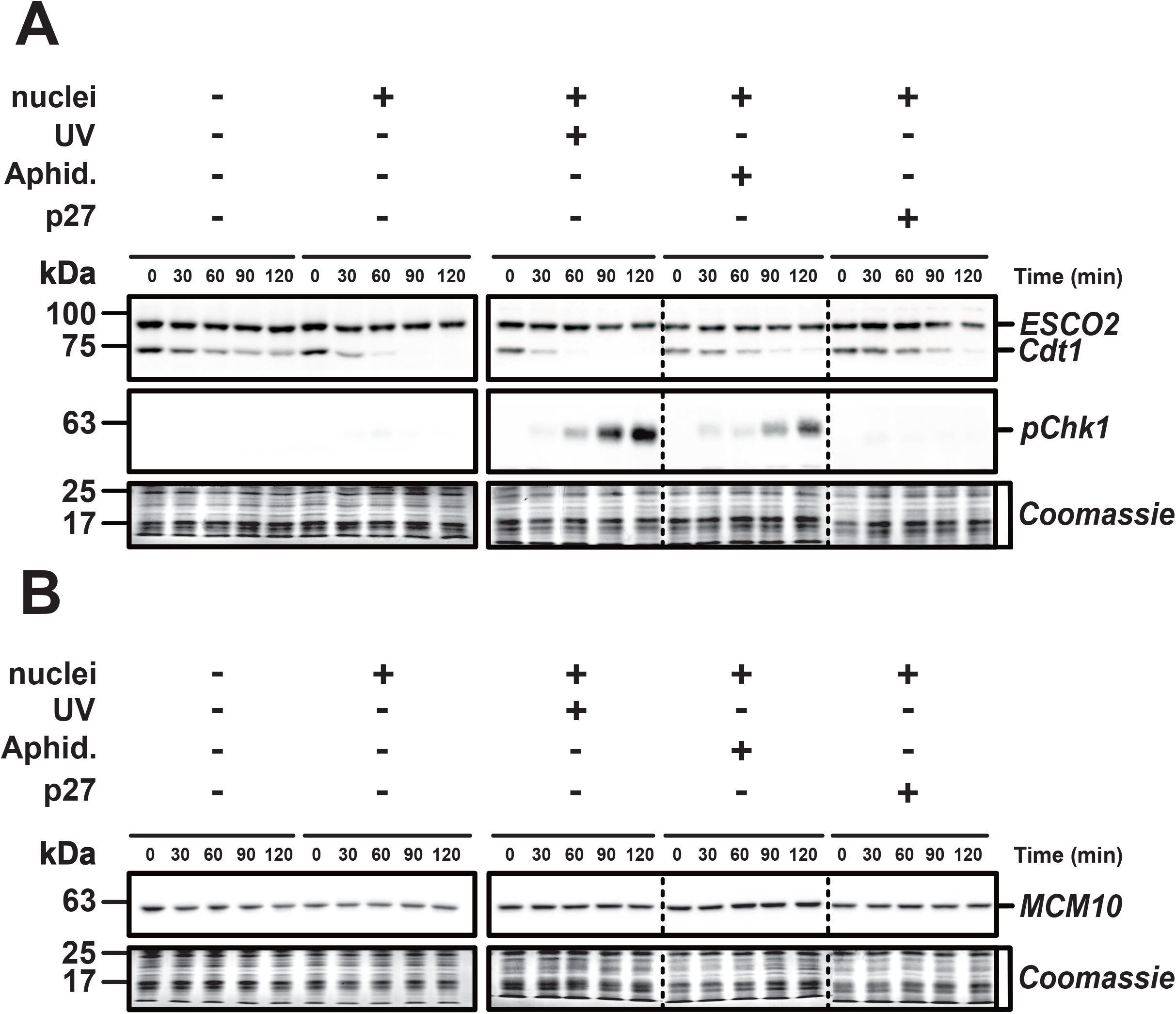
ESCO2 is stable during DNA damage signaling. **A**. Immunoblot analysis. Reactions were assembled as in Figure 1 with the indicated modifications. UV: sperm were UV treated before they were added to the extract. Aphid: the DNA replication inhibitor aphidicolin (150 μM final) was added. p27: Recombinant p27 protein was added to the extract before the addition of nuclei. Samples were collected at the indicated times and probed by immunoblot for the indicated proteins. Antibody specific for phosphorylated Chk1 kinase (pChk1) was used to confirm DNA damage signaling. **B.** Samples collected as in A were probed as indicated for MCM10. Solid outlines denote membrane fragments that were processed separately. Dotted lines denote where blot images were cropped.

Reliance on different specificity factors may explain the different sensitivity of ESCO2 and Cdt1 to the CUL4-dependent degradation machinery in egg extract. The CUL4 ubiquitin ligase can be activated by one of a number of DCAF subunits, which confer substrate specificity to the complex (13). Cdt1 is recognized by CUL4 when it is activated by DCAF2^Cdt2/Dtl^, and Cdt1 degradation requires its interaction with chromatin-bound PCNA, explaining the requirement for DNA replication (17). In contrast to Cdt1, ESCO2 has been proposed to be ubiquitinated by CUL4 activated by DCAF1^VprBP^, which is not well characterized in the *Xenopus* egg extract system and likely acts independently of PCNA (7). To test whether CUL4-DCAF1^VprBP^ is active in egg extracts, we tested the stability of the replication protein MCM10, which has been reported to be degraded through CUL4-DCAF1^VprBP^ in response to UV damage (27) (Fig. 3B). We found that endogenous MCM10 is stable in egg extracts, even in the presence of active DNA damage signaling. Proteomic analyses have suggested that the DCAF1^VprBP^ protein level is relatively constant during early *Xenopus* development (28) (Fig. S1). It is possible that DCAF1^Vprbp^-dependent degradation is not fully active until later in development, or has developmentally regulated changes in specificity.

Because we were unable to detect ESCO2 degradation during S phase, and because we could not with certainty identify appropriate control for DCAF1^Vprbp^-dependent degradation in the *Xenopus* system, we decided to investigate ESCO2 stability in somatic cells. To do this, we created stable HeLa cell lines in which the expression of a GFP-ESCO2 transgene is controlled by a tetracycline-inducible promoter. Asynchronously growing cells expressing GFP-ESCO2 were collected and analyzed by flow cytometry to assess ESCO2 levels relative to DNA content (Fig. 4A-B). In this asynchronous cell population, we found that ESCO2 levels were constant over the course of DNA replication with no statistically significant differences between cells in early or late S phase. Levels were significantly lower in G1 compared to G2/M (Fig. 4B) consistent with previous work showing that ESCO2 is targeted for degradation by APC^Cdh1^ (5). We conclude from this experiment that ESCO2 is stable during S phase and that APC-dependent modification in G1 likely accounts for all readily detectable ESCO2 degradation during cell cycle progression. We cannot rule out the possibility that the GFP tag or elevated ESCO2 expression level interfered with S phase mediated degradation of the transgene product.

**Figure 4.**
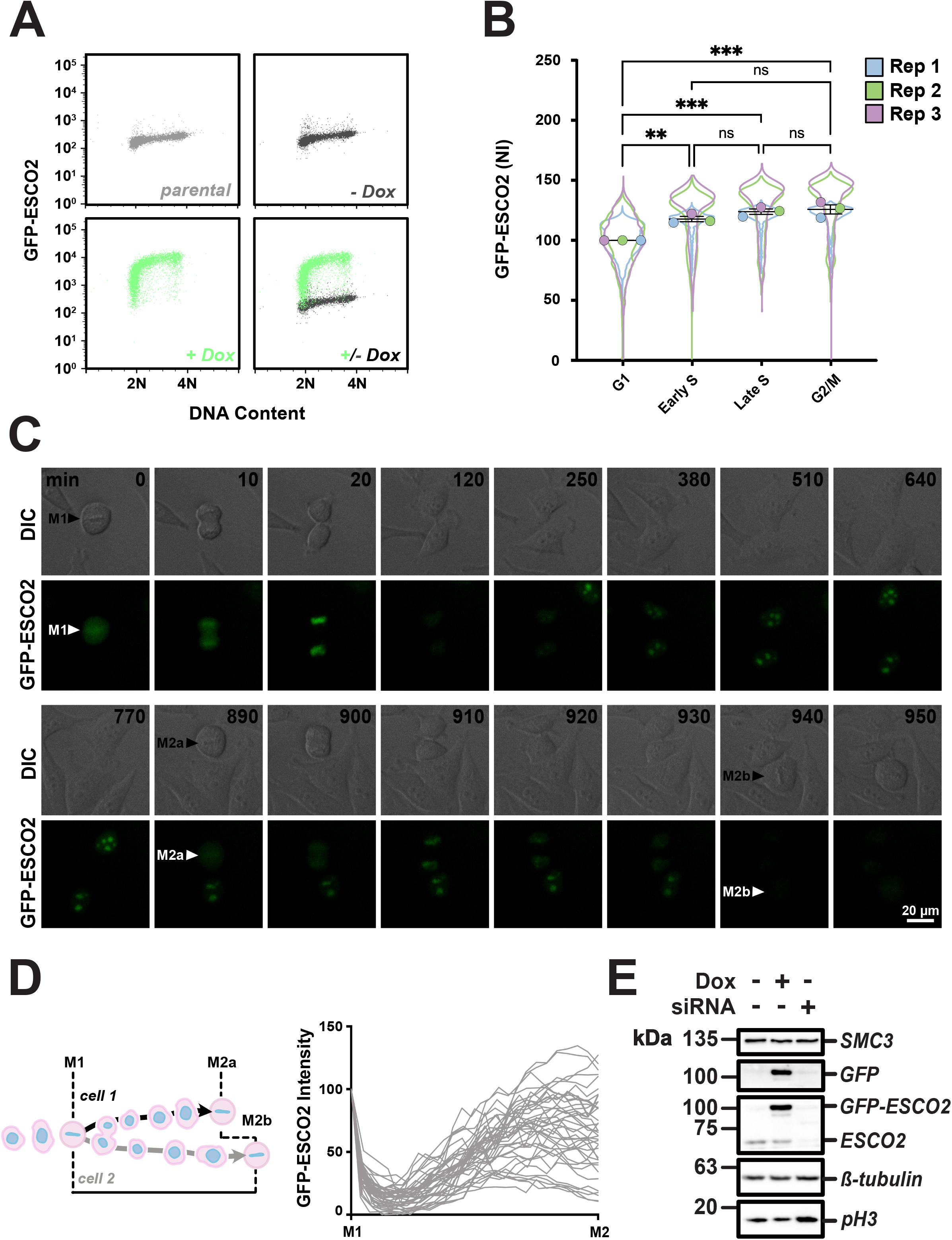
ESCO2 levels in cultured somatic cells. **A. Flow cytometry.** Live non-extracted HeLa cells expressing GFP-ESCO2 were collected and analyzed for GFP levels and DNA content. Parental and uninduced GFP-ESCO2 (no doxycycine added) cell lines are also shown. n≥ 5500. **B. Quantification of GFP-ESCO2 levels during cell cycle progression.** The flow cytometry experiment shown in A, was repeated three times and are shown together as a SuperPlot (44). Mean values from each replicate, normalized to their G1 mean value, are plotted together. NI = Normalized intensity. Error bars = SEM. ns = not significant, p > 0.05; **, p < 0.01; ***, p < 0.001; ****, p < 0.0001. **C. Time lapse imaging.** Shown are selected frames from time-lapse imaging of HeLa cells expressing GFP-ESCO2. Intervals were chosen to highlight details of cell division. Time elapsed (in minutes) since metaphase is indicated by numbers in black font. The complete movie is shown in Video S1. **D. Quantification of GFP signal.** Schematic at left shows how GFP fluorescence intensity was scored across a complete cell division cycle, from an initial metaphase (M1) to metaphase 2 (M2) for each daughter cell, and graphed in aggregate at right. n = 20 M1 cells, 40 M2 cells. cell analyzed. **E. Immunoblot analysis.** The cell line used in A-D was treated either with doxycycline to induce transgene expression, or siRNA to deplete the endogenous protein, and cell lysates were analyzed by immunoblot for the indicated proteins. Solid outlines denote membrane fragments that were processed separately. SMC3, a subunit of cohesin, and *β*-tubulin served as a loading controls. A replicate blot was prepared and probed for GFP. Phosphorylated H3 (pH3), a marker of mitotic cells, confirmed increased mitotic index in cells depleted of ESCO2 seen previously (42).

To further characterize ESCO2 stability during cell cycle progression, we analyzed our GFP-ESCO2 cell line by time-lapse microscopy under inducing conditions (Fig. 4C, Video S1, Video S2). As expected, the GFP signal was nuclear in interphase and dispersed from chromatin as cells entered M phase (4). Consistent with our previous results, GFP-ESCO2 also accumulated in nucleoli (29). This may be due in part to overexpression, or to the presence of the GFP tag which is known to partition to this location (30). To quantify ESCO2 turnover, we measured the total cellular GFP signal as single cells progressed from metaphase to the subsequent metaphase of each resulting daughter cell (Fig. 4D). Consistent with the flow cytometry data, the GFP-ESCO2 signal was high in metaphase, and dropped after the cell division, with minimal levels attained approximately 160 minutes after anaphase. As cells further progressed through the cell cycle, the GFP-ESCO2 signal rose and remained high until the next metaphase. We saw no clear evidence of ESCO2 loss after the initial drop at anaphase (Fig. 4F). Thus by both by flow cytometry and image analysis, we saw no evidence of a decrease in ESCO2 accompanying DNA replication. We cannot rule out the possibility that overexpression affected turnover at this time, but note that the predicted APC-dependent loss of ESCO2 in G1 was readily detected. Immunoblot analysis indicated that in our cell line GFP-ESCO2 was expressed at ~6.5 times the level of endogenous ESCO2 (Fig. 4E), with some variability between cells (Video S2). We conclude from these experiments that ESCO2 is largely stable during S phase in somatic cells.

## Discussion

We have tested ESCO2 stability during DNA replication, both by assessing the endogenous protein in *Xenopus* egg extract and by using a stably-expressed fusion protein in cultured somatic cells. Although it has been proposed that ESCO2 is degraded by CUL4-dependent mechanisms during or shortly after S phase, here we find that ESCO2 is stable during DNA replication in both systems.

There are several possible explanations for the different conclusions drawn here and those made previously. Previous work showed that ESCO2 levels were elevated (compared to controls) in G2 cells following depletion of DCAF1^VprBP^, but ESCO2 was not shown to be directly modified by the CUL4 ubiquitin ligase(7). The increase in ESCO2 in the absence of DCAF1^VprBP^ may be an indirect result of the impact of DCAF1^VprBP^ on cell cycle progression. DCAF1^VprBP^ was originally characterized based on its interaction with lentiviral R proteins, which is thought to alter cell cycle progression, although the underlying mechanism. DCAF1^VprBP^ has cell cycle impacts independent of viral infection (31). In unperturbed cells, the tumor suppressor NF2^Merlin^, an ezrin-moesin-radixin (ERM) family protein that is mutated in neurofibromatosis type 2, binds to and inhibits the nuclear Cul4-DCAF1^VprBP^ complex (32–34). Depletion of CUL4-DCAF1^VprBP^ phenocopies the loss of Merlin function, suggesting that CUL4-DCAF1^VprBP^ normally has an antiproliferative function. In other models, such as T cell development, DCAF1^VprBP^ is required for cell cycle entry (35). The ubiquitination targets of this complex that modulate proliferative activity have not been identified, although certain transcription factors may be important (36). DCAF1^VprBP^ has non-proteolytic impacts on several proteins, including the p53 tumor suppressor, the FoxM1 transcription factor, and the SAMHD1 viral restriction factor (37). The myriad activities of the CUL4-DCAF1^VprBP^ complex make understanding its precise contribution to ESCO2 stability unclear. Fully assessing the function and activity of CUL4-DCAF1 in embryonic extracts is an important topic beyond the scope of this current work.

We initially set out in this work to identify degrons in the ESCO2 protein that might promote protein turnover during S phase, thereby controlling cohesin in G2 cells. We have found the ESCO2 protein to be remarkably stable in egg extract, and in somatic cells only obviously reduced in G1, most likely through the previously-reported APC degron in the ESCO2 N terminus (5). It remains possible that S phase degradation of ESCO2 is developmentally regulated or cell-type specific, perhaps occurring through mechanisms that are not yet established in the early frog embryo or elaborated later in development. However, we found no evidence for significant loss of ESCO2 during DNA replication in unperturbed somatic cells or embryonic nuclei. Although previous work makes clear that ESCO2 makes complex interactions with the DNA replication machinery (6, 9, 10, 38), whether ESCO2 has functions after DNA replication remains unclear at this time.

## Experimental procedures

### Gels and Immunoblots

Egg extract samples were diluted in sample buffer (1:10) and loaded onto a 7-15% gradient SDS-PAGE gel to resolve proteins. Gels were cut near the 25 kDa marker and the lower fragment was stained with a colloidal Coomassie stain. Proteins were transferred from the remainder of the gel to nitrocellulose membrane using the Trans-Blot Turbo (Bio-Rad). Membranes were incubated in a 5% milk in 1X Tris-buffered saline, 0.05% Tween 20 (TBST) for 45 minutes at room temperature, probed using primary antibodies overnight at 4°C, washed three times in 1X TBST, probed with horseradish peroxidase (HRP)-conjugated secondary antibodies for 45 minutes at room temperature, washed three times in 1X TBST, and one time with 1X TBS. Signals were detected with chemiluminescent substrate (Licor Biosciences) and imaged with the Azure C600 CCD imager (Azure Biosystems). Intensity measurements were made using Image Studio Lite (Licor Biosciences) using median background subtraction and top/bottom setting.

### Antibodies

Primary antibodies to P-Chk1 (Cell Signaling, rabbit, 1:1200), ß-tubulin (DSHB, mouse, 1:2000), hESCO2 (Bethyl Labs, rabbit,1:3000), GFP (Millipore Sigma, mouse, 1:5000), and hPH3 (Upstate, rabbit, 1:1000) were obtained commercially. Antibodies to xlCdt1 (rabbit, 1:2000) and MCM10 (rabbit, 1:5000) were kindly provided by Johannes Walter(14, 39) and xlESCO2 (rabbit, 1:2000) and hSmc3 (rabbit, 1:1000) were generated by the Rankin lab (2). P-Chk1 antibodies were diluted in 5% BSA blocking solution (BSA in 1X TBST) with 0.1% NaN_3_ and all others were diluted in 5% milk blocking solution with 0.1% NaN_3_. Goat anti-rabbit (Life Technologies, 1:10,000) and donkey anti-mouse secondary (Jackson ImmunoResearch Laboratories, 1:5000) antibodies were used diluted in a 5% milk blocking solution.

### Xenopus egg extracts

All animal work in this study has been done with the approval and oversight of the OMRF Institutional Animal Care and Use Committee. Egg extracts were prepared according to established protocols(40). *Xenopus laevis* eggs were collected in 1X MMR (5 mM HEPES, pH 7.7; 100 mM NaCl; 2 mM KCl; 1 mM MgCl_2_; 2 mM CaCl_2_; 0.1 mM EDTA), dejellied in a 2% cysteine in water containing 1mM EGTA, and washed 4-5 times with XBE2 (2 M KCl; 1 M MgCl_2_; 1 M CaCl_2_; 1.7% sucrose; 0.5 M K-EGTA; 1M HEPES, pH 7.7). Eggs were then transferred to fill 14 X 89 mm round-bottom tubes (Beckman Coulter, 331372) supplemented with protease inhibitors (10 μg/ml each of leupeptin, pepstatin, and chymostatin, final). The eggs were then packed by centrifugation (1,000 RPM for 1 minute at 4 °C) using a Beckman JS13.1 rotor. Excess buffer was removed from packed eggs which were then crushed by centrifugation at 10,000 RPM for 10 minutes at 4 °C in the same rotor. The cytosolic layer was removed via side puncture 16-G needle, and supplemented with protease inhibitors (10 μg/ml each of leupeptin, pepstatin, and chymostatin), cytochalasin B (10 μg/ml), 15% LFB1/50 (40 mM HEPES, pH 8.0; 20 mM K_2_HPO_4_/KH_2_PO_4_, pH 8.0; 2 mM MgCl_2_; 1mM EGTA; 2 mM DTT; 10% sucrose; plus 50 mM KCl), transferred to round-bottom tubes (Beckman Coulter, 344057), and spun at 30,000 RPM in a Beckman Sw55 rotor for 20 minutes at 4 °C. The lipid plug was pushed aside and the cytosol, including floating membrane layer, was collected using a pipette tip. The extract was supplemented with 2% v:v glycerol and snap-frozen in 100 μl aliquots using liquid nitrogen and stored at −80 °C.

Prior to use, extracts were quickly thawed in hand, set on ice, and supplemented with freshly prepared 35X stock of energy mix (20 mM phosphocreatine, 5 μg/ml creatine phosphokinase, and 2 mM ATP) made from frozen components. To release extract from CSF arrest, a freshly made 10 mM CaCl_2_ solution was added (0.4 mM final in extract) and extracts were incubated in a 20°C water bath for 30 minutes. Where appropriate, cycloheximide (Sigma Aldrich), was added to CSF extracts at 10 mg/ml (250 μg/ml final), before the addition of CaCl_2_. Aphidicolin (VWR) treatment was (100 μg/ml final) from a 10mg/ml stock solution following CSF release and before sperm addition. Purified H6-p27 protein at 1 mg/ml was added to egg extracts (20 μg/ml final) following CSF release and before sperm addition.

### Sperm nuclei

Preparation of demembranated sperm nuclei was as previously described (41). Briefly, freshly isolated testes were minced and washed in buffer X (100 mM HEPES pH 7.5, 800 mM KCl, 150 mM NaCl, 50 mM MgCl_2_, 10 mM EDTA, 200 mM sucrose), vortexed, and spun with mild centrifugation (10 seconds at 1,000 RPM), repeating until the supernatant was clear. Supernatants were combined and centrifuged twice (50 seconds at 1,500 RPM and 10 minutes at 4,000 RPM at 4°C). The pellet was then resuspended in buffer X and layered on a sucrose gradient and centrifuged 25 minutes at 33,000 RPM at 4°C. The sperm pellet was again resuspended in buffer X and centrifuged (10 minutes at 5,000 RPM at 4 °C), resuspended in buffer X mix #1 (buffer X, 0.4% Triton X-100, and LPC), and incubated while rotating at 4°C for 30 minutes. The resulting solution is layered over a sucrose step gradient and centrifuged (10 minutes at 2,100 RPM at room temperature), resuspended in buffer X mix #2 (buffer X, 3% BSA and LPC), and centrifuged (10 minutes at 2,100 RPM at room temperature) twice, and resuspended in a final buffer X mix (buffer X, 3% BSA, LPC, and 1 mM DTT). Sperm concentration was determined using a hemocytometer. Final sperm preparations were frozen in 10 μl aliquots and stored at −80 °C.

To damage sperm nuclei, a 5 μl drop of ice-cold sperm (1.2×10^5^/ μl) was deposited on parafilm at room temperature and irradiated in a Stratalinker 1800 (Stratagene) with ~700 μJ/m^2^ UV. Control sperm were left on the bench on parafilm at room temperature for 5 minutes.

### Cell culture

Cells were routinely cultured in Dulbecco’s modified Eagle’s medium (DMEM, Corning) supplemented with 10% fetal bovine serum (FBS, R&D Systems), and maintained at 37°C in 5% CO_2_. Cell lines with a doxycycline-inducible NLS-GFP-tagged ESCO2 transgene were generated using HeLa Flp-In T-Rex ESCO1 KO cells as previously (42). Cells were selected using 200 μg/ml hygromycin B (Gold Biotechnology), and single colonies were isolated using trypsin-soaked filter paper. Transgene expression was induced with 24-hour incubation in media supplemented with 2 μg/ml doxycycline (VWR). SiRNA-mediated depletion of endogenous ESCO2 was done with 20 nM siRNA (Dharmacon, J-025788-09, target: CGAGUGAUCUAUAAGCCAA) using Lipofectamine RNAiMAX (Invitrogen) according to the manufacturer’s instructions, in Opt-MEM serum free medium (Invitrogen). Following 12 hours in transfection mix, the media was replaced with fresh standard medium supplemented with 2 μg/ml of doxycycline, cells were incubated for an additional 24 hours, and then processed for immunoblot.

### Flow cytometry

To analyze live HeLa cells by flow cytometry Hoechst 33342 (4 μg/ml) was added to the media for 45 minutes prior to harvesting. The media was collected and the cells were washed with PBS + 4 μg/ml Hoechst, harvested with trypsin, and resuspended in reserved media. The cells were centrifuged at 1500 RPM for 5 minutes, resuspended in PBS + 4 μg/ml Hoechst 33342, and run on a FACSCelesta flow cytometer (BD Biosciences). Samples were analyzed using FlowJo v10.5.3 (TreeStar). Single cells were identified using the forward and side scatter (Fig. S2A), and the Cell Cycle univariate modeling tool (FlowJo, Watson Pragmatic algorithm) based on DNA content was used to assign cell cycle phases to each dataset (Fig. S2B). S phase was further subdivided into early and late populations by imposing a gate at the midpoint of DNA content within the S phase group (Fig. S2C). The fluorescence intensity values of each phase were exported into Prism and normalized to the average G1 intensity for each of the three biological replicates.

### Live-cell imaging

Cells induced to express GFP-ESCO2 overnight were imaged in Opti-MEM (Gibco) supplemented with 2 μg/ml of Doxycycline and 250 nM siR-DNA at 37°C in a 5% CO_2_ atmosphere using a stage top incubator (Tokai Hit). Images were collected every 10 minutes for 48 hours using a 20X S Plan Fluor ELWD objective lens on a Nikon Eclipse TE2000-E equipped with the Perfect Focus and triggered acquisition, with a Hamamatsu Orca-Flash4.0 CMOS camera and Lumencor light engine light source. Images were analyzed using NIS Elements software. Beginning with metaphase figures, the background-subtracted sum cellular GFP intensity of each of 20 cells was measured every 40 minutes until the subsequent metaphase of the two daughter cells and plotted, each normalized to the signal intensity of the maternal cell. The time (x) axis for each cell was normalized to individual cell cycle lengths, so that measurements at M1 and M2 for all cells are aligned for direct comparison.

### Statistical analysis

Prism v9.3 (Graph-Pad Software) was used to plot data and perform statistical analysis. For analyses with multiple comparisons, we used an ordinary one-way ANOVA and Tukey’s multiple comparison test, with a single pooled variance. For analysis of flow cytometry data, the fluorescence intensity values were exported into Prism and normalized to the average G1 intensity for each of the three biological replicates, which were then overlaid in a Superplot (43).

## Supporting information

Supplemental figure legends.docx

Video S1.avi

Video S2.avi

## Data availability

Data from this study are included in this manuscript and associated supplemental files. Any additional details gladly provided upon request.

## Acknowledgments

We are grateful to Johannes Walter for contributing the anti-Cdt1 and anti-MCM10 antibodies used in this study. We also thank the members of the Rankin lab for thoughtful discussion in preparation for this manuscript and to all members of the Program in Cell Cycle and Cancer Biology for their input and discussions during this work.

## Author contributions

S.R. conceptualization; A.J., B.R., S.R. formal analysis; S.R. funding acquisition; A.J., B.R. investigation; A.J., B.R., S.R. methodology; S.R. project administration; J.C., S.R. resources; S.R. supervision; A.J., B.R., S.R., visualization; A.J., B.R., S.R. writing – original draft; A.J., B.R., S.R. writing - review & editing.

## Funding and additional information

This study was supported by National Institutes of Health grant R01GM101250 to S.R. The content is solely the responsibility of the authors and does not necessarily represent the official views of the National Institutes of Health.

## Abbreviations

APC: (Anaphase promoting complex)
Cdh1: (Cdc20 homolog 1)
Cdt1: (Chromatin licensing and DNA replication factor 1)
Chk1: (Checkpoint kinase 1)
CRL4: (Cullin-RING ubiquitin ligase complex 4)
CSF: (Cytostatic factor)
Cul4: (Cullin-4)
DCAF1: (DDB1-Cul4-associating factor 1)
DDB1: (DNA damage-binding protein 1)
Dox: (doxycycline)
Eco1: (Establishment of cohesion 1)
ESCO2: (Establishment of Cohesion 1 homolog 2)
FoxM1: (Forkhead box M1)
MCM: (Minichromosome maintenance)
PCNA: (Proliferating cell nuclear antigen)
PIP: (PCNA-interacting protein)
SAMHD1: (SAM and HD domain containing deoxynucleoside triphosphate triphosphohydrolase 1)
SMC3: (Structural Maintenance of Chromosomes 3)
VprBP: (Viral protein R binding protein)
WAPL: (Wings apart-like protein homolog)

---

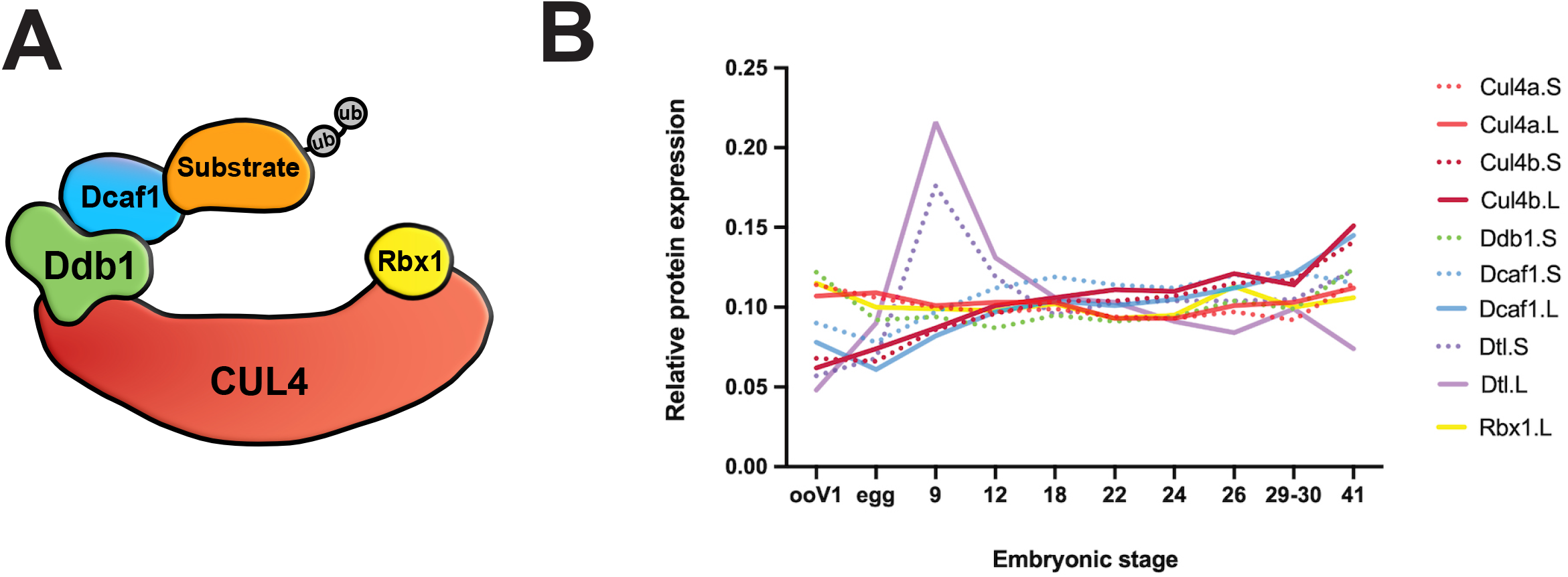

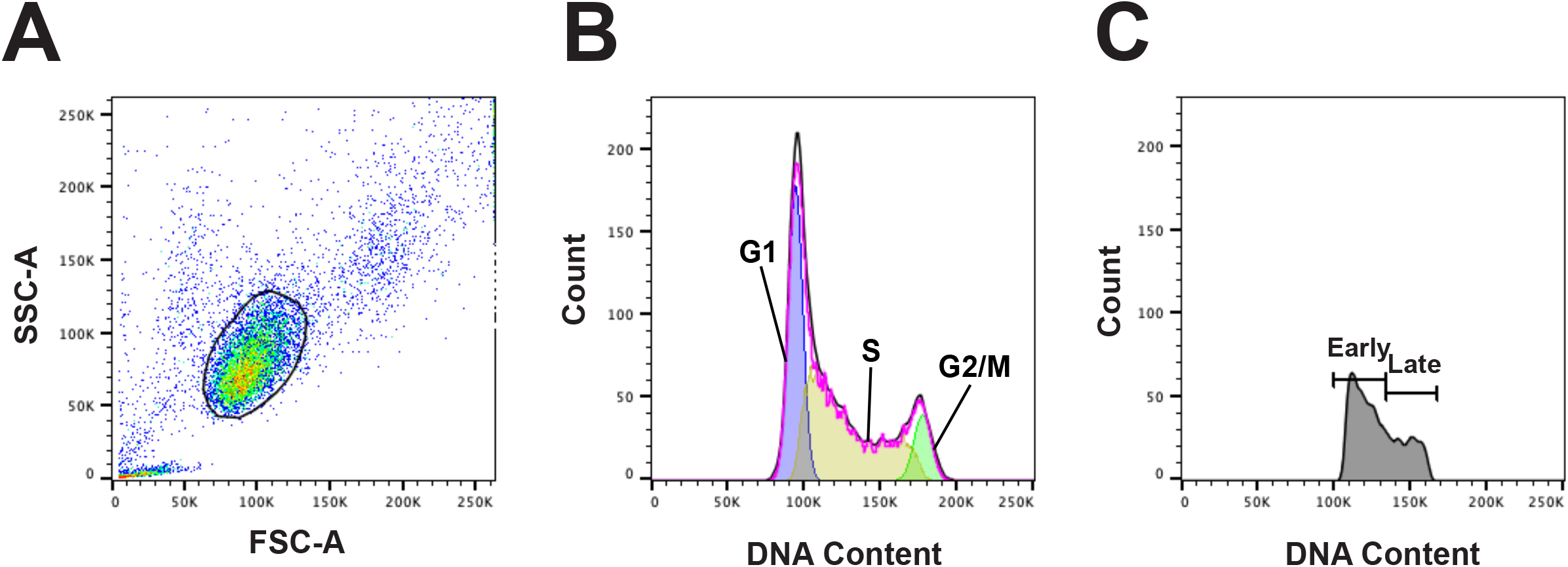

## References cited

1. Unal, E., Heidinger-Pauli, J. M., Kim, W., Guacci, V., Onn, I., Gygi, S. P., and Koshland, D. E. (2008) A molecular determinant for the establishment of sister chromatid cohesion. Science. 321, 566–569

2. Zhang, J., Shi, X., Li, Y., Kim, B.-J., Jia, J., Huang, Z., Yang, T., Fu, X., Jung, S. Y., Wang, Y., Zhang, P., Kim, S.-T., Pan, X., and Qin, J. (2008) Acetylation of Smc3 by Eco1 Is Required for S Phase Sister Chromatid Cohesion in Both Human and Yeast. Molecular Cell. 31, 143–151

3. Sutani, T., Kawaguchi, T., Kanno, R., Itoh, T., and Shirahige, K. (2009) Budding yeast Wpl1(Rad61)-Pds5 complex counteracts sister chromatid cohesion-establishing reaction. Current biology□: CB. 19, 492–497

4. Hou, F., and Zou, H. (2005) Two human orthologues of Eco1/Ctf7 acetyltransferases are both required for proper sister-chromatid cohesion. Molecular biology of the cell. 16, 3908–3918

5. Lafont, A. L., Song, J., and Rankin, S. (2010) Sororin cooperates with the acetyltransferase Eco2 to ensure DNA replication-dependent sister chromatid cohesion. Proceedings of the National Academy of Sciences of the United States of America. 107, 20364–20369

6. Song, J., Lafont, A., Chen, J., Wu, F. M., Shirahige, K., and Rankin, S. (2012) Cohesin acetylation promotes sister chromatid cohesion only in association with the replication machinery. Journal of Biological Chemistry. 287, 34325–34336

7. Minamino, M., Tei, S., Negishi, L., Kanemaki, M. T., Yoshimura, A., Sutani, T., Bando, M., and Shirahige, K. (2018) Temporal Regulation of ESCO2 Degradation by the MCM Complex, the CUL4-DDB1-VPRBP Complex, and the Anaphase-Promoting Complex. Current biology: CB. 28, 2665–2672.e5

8. Lelij, P. van der, Godthelp, B. C., Zon, W. van, Gosliga, D. V., Oostra, A. B., Steltenpool, J., Groot, J. de, Scheper, R. J., Wolthuis, R. M., Waisfisz, Q., Darroudi, F., Joenje, H., and Winter, J. P. de (2009) The Cellular Phenotype of Roberts Syndrome Fibroblasts as Revealed by Ectopic Expression of ESCO2. PloS one. 4, e6936

9. Ivanov, M. P., Ladurner, R., Poser, I., Beveridge, R., Rampler, E., Hudecz, O., Novatchkova, M., Hériché, J.-K., Wutz, G., Lelij, P. van der, Kreidl, E., Hutchins, J. R., Ekker, H. A., Ellenberg, J., Hyman, A. A., Mechtler, K., and Peters, J.-M. (2018) The replicative helicase MCM recruits cohesin acetyltransferase ESCO2 to mediate centromeric sister chromatid cohesion. The EMBO Journal. 37, e97150

10. Bender, D., Silva, E. M. L. D., Chen, J., Poss, A., Gawey, L., Rulon, Z., and Rankin, S. (2019) Multivalent interaction of ESCO2 with the replication machinery is required for sister chromatid cohesion in vertebrates. Proceedings of the National Academy of Sciences of the United States of America. 107, 201911936

11. Davey, N. E., and Morgan, D. O. (2016) Building a Regulatory Network with Short Linear Sequence Motifs: Lessons from the Degrons of the Anaphase-Promoting Complex. Molecular Cell. 64, 12–23

12. Visintin, R., Prinz, S., and Amon, A. (1997) CDC20 and CDH1: a family of substrate-specific activators of APC-dependent proteolysis. Science. 278, 460–463

13. Jin, J., Arias, E. E., Chen, J., Harper, J. W., and Walter, J. C. (2006) A family of diverse Cul4-Ddb1-interacting proteins includes Cdt2, which is required for S phase destruction of the replication factor Cdt1. Molecular Cell. 23, 709–721

14. Arias, E. E., and Walter, J. C. (2005) Replication-dependent destruction of Cdt1 limits DNA replication to a single round per cell cycle in Xenopus egg extracts. Genes & Development. 19, 114–126

15. Arias, E. E., and Walter, J. C. (2006) PCNA functions as a molecular platform to trigger Cdt1 destruction and prevent re-replication. Nature Cell Biology. 8, 84–90

16. Havens, C. G., Shobnam, N., Guarino, E., Centore, R. C., Zou, L., Kearsey, S. E., and Walter, J. C. (2012) Direct role for proliferating cell nuclear antigen in substrate recognition by the E3 ubiquitin ligase CRL4Cdt2. Journal of Biological Chemistry. 287, 11410–11421

17. Havens, C. G., and Walter, J. C. (2009) Docking of a specialized PIP Box onto chromatin-bound PCNA creates a degron for the ubiquitin ligase CRL4Cdt2. Molecular Cell. 35, 93–104

18. Jevitt, A. M., and Rankin, S. (2022) Xenopus: From Basic Biology to Disease Models in the Genomic Era (Fainsod, A., and Moody, S. A. eds), 10.1201/9781003050230-3

19. Rankin, S. (2019) Reconstituting Nuclear and Chromosome Dynamics Using Xenopus Extracts., p. pdb.top097105, Cold Spring Harbor Protocols, 2019, pdb.top097105

20. Newport, J., and Kirschner, M. (1982) A major developmental transition in early Xenopus embryos: II. Control of the onset of transcription. Cell. 30, 687–696

21. Lyons, N. A., and Morgan, D. O. (2011) Cdk1-dependent destruction of Eco1 prevents cohesion establishment after S phase. Molecular Cell. 42, 378–389

22. Byun, T. S., Pacek, M., Yee, M.-C., Walter, J. C., and Cimprich, K. A. (2005) Functional uncoupling of MCM helicase and DNA polymerase activities activates the ATR-dependent checkpoint. Genes & Development. 19, 1040–1052

23. Hekmat-Nejad, M., You, Z., Yee, M. C., Newport, J. W., and Cimprich, K. A. (2000) Xenopus ATR is a replication-dependent chromatin-binding protein required for the DNA replication checkpoint. Current Biology. 10, 1565–1573

24. Recolin, B., Laan, S. V. D., and Maiorano, D. (2012) Role of replication protein A as sensor in activation of the S-phase checkpoint in Xenopus egg extracts. Nucleic Acids Res. 40, 3431–3442

25. Kumagai, A., Yakowec, P. S., and Dunphy, W. G. (1998) 14-3-3 Proteins Act as Negative Regulators of the Mitotic Inducer Cdc25 in Xenopus Egg Extracts. Mol Biol Cell. 9, 345–354

26. Walworth, N. C., and Bernards, R. (1996) rad-Dependent Response of the chk1-Encoded Protein Kinase at the DNA Damage Checkpoint. Science. 271, 353–356

27. Kaur, M., Khan, Md. M., Kar, A., Sharma, A., and Saxena, S. (2012) CRL4-DDB1-VPRBP ubiquitin ligase mediates the stress triggered proteolysis of Mcm10. Nucleic Acids Res. 40, 7332–7346

28. Peshkin, L., Lukyanov, A., Kalocsay, M., Gage, R. M., Wang, D., Pells, T. J., Karimi, K., Vize, P. D., Wühr, M., and Kirschner, M. W. (2019) The protein repertoire in early vertebrate embryogenesis. Biorxiv. 10.1101/571174

29. Bender, D., Silva, E. M. L. D., Chen, J., Poss, A., Gawey, L., Rulon, Z., and Rankin, S. (2020) Multivalent interaction of ESCO2 with the replication machinery is required for sister chromatid cohesion in vertebrates. Proc National Acad Sci. 117, 1081–1089

30. Martin, R. M., Ter-Avetisyan, G., Herce, H. D., Ludwig, A. K., Lättig-Tünnemann, G., and Cardoso, M. C. (2015) Principles of protein targeting to the nucleolus. Nucleus. 6, 314–325

31. Han, X.-R., Sasaki, N., Jackson, S. C., Wang, P., Li, Z., Smith, M. D., Xie, L., Chen, X., Zhang, Y., Marzluff, W. F., and Xiong, Y. (2020) CRL4DCAF1/VprBP E3 ubiquitin ligase controls ribosome biogenesis, cell proliferation, and development. Sci Adv. 6, eabd6078

32. Trofatter, J. A., MacCollin, M. M., Rutter, J. L., Murrell, J. R., Duyao, M. P., Parry, D. M., Eldridge, R., Kley, N., Menon, A. G., Pulaski, K., Haase, V. H., Ambrose, C. M., Munroe, D., Bove, C., Haines, J. L., Martuza, R. L., MacDonald, M. E., Seizinger, B. R., Short, M. P., Buckler, A. J., and Gusella, J. F. (1993) A novel moesin-, ezrin-, radixin-like gene is a candidate for the neurofibromatosis 2 tumor suppressor. Cell. 72, 791–800

33. Rouleau, G. A., Merel, P., Lutchman, M., Sanson, M., Zucman, J., Marineau, C., Hoang-Xuan, K., Demczuk, S., Desmaze, C., Plougastel, B., Pulst, S. M., Lenoir, G., Bijlsma, E., Fashold, R., Dumanski, J., Jong, P. de, Parry, D., Eldrige, R., Aurias, A., Delattre, O., and Thomas, G. (1993) Alteration in a new gene encoding a putative membrane-organizing protein causes neuro-fibromatosis type 2. Nature. 363, 515–521

34. Cooper, J., Li, W., You, L., Schiavon, G., Pepe-Caprio, A., Zhou, L., Ishii, R., Giovannini, M., Hanemann, C. O., Long, S. B., Erdjument-Bromage, H., Zhou, P., Tempst, P., and Giancotti, F. G. (2011) Merlin/NF2 Functions Upstream of the Nuclear E3 Ubiquitin Ligase CRL4DCAF1 to Suppress Oncogenic Gene Expression. Sci Signal. 4, pt6

35. Guo, Z., Kong, Q., Liu, C., Zhang, S., Zou, L., Yan, F., Whitmire, J. K., Xiong, Y., Chen, X., and Wan, Y. Y. (2016) DCAF1 controls T-cell function via p53-dependent and -independent mechanisms. Nat Commun. 7, 10307

36. Wang, X., Arceci, A., Bird, K., Mills, C. A., Choudhury, R., Kernan, J. L., Zhou, C., Bae-Jump, V., Bowers, A., and Emanuele, M. J. (2017) VprBP/DCAF1 Regulates the Degradation and Nonproteolytic Activation of the Cell Cycle Transcription Factor FoxM1. Mol Cell Biol. 10.1128/mcb.00609-16

37. Nakagawa, T., Mondal, K., and Swanson, P. C. (2013) VprBP (DCAF1): a promiscuous substrate recognition subunit that incorporates into both RING-family CRL4 and HECT-family EDD/UBR5 E3 ubiquitin ligases. Bmc Mol Biol. 14, 22–22

38. Higashi, T. L., Ikeda, M., Tanaka, H., Nakagawa, T., Bando, M., Shirahige, K., Kubota, Y., Takisawa, H., Masukata, H., and Takahashi, T. S. (2012) The prereplication complex recruits XEco2 to chromatin to promote cohesin acetylation in Xenopus egg extracts. Current biology: CB. 22, 977–988

39. Wohlschlegel, J. A., Dhar, S. K., Prokhorova, T. A., Dutta, A., and Walter, J. C. (2002) Xenopus Mcm10 Binds to Origins of DNA Replication after Mcm2-7 and Stimulates Origin Binding of Cdc45. Mol Cell. 9, 233–240

40. Gillespie, P. J., Gambus, A., and Blow, J. J. (2012) Preparation and use of Xenopus egg extracts to study DNA replication and chromatin associated proteins. Methods (San Diego, Calif). 57, 203–213

41. Chan, R. C., and Forbes, D. J. (2006) Xenopus Protocols, Cell Biology and Signal Transduction. Methods Mol Biology. 322, 289–300

42. Alomer, R. M., Silva, E. M. L. da, Chen, J., Piekarz, K. M., McDonald, K., Sansam, C. G., Sansam, C. L., and Rankin, S. (2017) Esco1 and Esco2 regulate distinct cohesin functions during cell cycle progression. Proceedings of the National Academy of Sciences of the United States of America. 114, 9906–9911

43. Lord, S. J., Velle, K. B., Mullins, R. D., and Fritz-Laylin, L. K. (2020) SuperPlots: Communicating reproducibility and variability in cell biology. J Cell Biol. 219, 94–10

44. Lord, S. J., Velle, K. B., Mullins, R. D., and Fritz-Laylin, L. K. (2020) SuperPlots: Communicating reproducibility and variability in cell biology. J Cell Biology. 219, e202001064

