## Supplemental figure legends.docx for "The cohesin modifier ESCO2 is stable during DNA replication"

**Supporting information**

**Figure S1.** **Relative protein expression during early development**. **A.** Illustration of proteins that comprise the Cul4-DCAF1^VprBP^ complex. **B.** Plot of relative protein expression for each component shown in B*.* For the genes that have expression from homeologous copies in the allotetraploid *Xenopus laevis*, the shorter and longer homeologs are signified with .S or .L respectively. Colors are coordinated with illustration in A. This graph is provided to illustrate developmental changes in the levels of each protein, rather than for direct comparison between proteins, which may not be valid.

**Figure S2.** Gating strategy and cell cycle assignment for flow cytometry analysis. **A.** Selection of single-cell population to exclude debris and cell clumps. SSC-A = side scatter area, FSC-A = forward scatter area. **B.** Cell cycle distribution of the single-cell population selected in A following analysis with the FlowJo Cell Cycle tool. **C.** S phase cells identified in B were bisected into early and late S phase at the midpoint of DNA content.

**Video S1:** Time-lapse movie of HeLa GFP-ESCO2 expressing cell from which stills shown in Figure 4 were extracted. Differential Interference Contrast (DIC) left and GFP (right). Elapsed time (hours: minutes: seconds) is shown at the top left.

**Video S2:** Time-lapse imaging of GFP-ESCO2 expressing cells, large field of view. Differential Interference Contrast (DIC) (left) and GFP (right). White arrow heads indicate examples of metaphase cells that undergo cell division over the course of the movie. Elapsed time (hours: minutes: seconds) is shown at top left.
